# Bisphenol A promotes stress granule assembly and modulates the integrated stress response

**DOI:** 10.1101/673194

**Authors:** Marta M. Fay, Daniella Columbo, Cecelia Cotter, Chandler Friend, Shawna Henry, Megan Hoppe, Paulina Karabelas, Corbyn Lamy, Miranda Lawell, Sarah Monteith, Christina Noyes, Paige Salerno, Jingyi Wu, Hedan Mindy Zhang, Paul J. Anderson, Nancy Kedersha, Pavel Ivanov, Natalie G. Farny

**Affiliations:** Division of Rheumatology, Immunology, and Allergy, Brigham and Women’s Hospital, Boston, USA; Department of Medicine, Harvard Medical School, Boston, MA, USA; Department of Biology and Biotechnology, Worcester Polytechnic Institute, Worcester, MA, USA; Broad Institute of Harvard and MIT, Cambridge, MA, USA; Currently at Gonzaga University, Spokane, WA, USA

**Author notes:** Co-corresponding authors: Pavel Ivanov, Natalie G. Farny, Department of Biology and Biotechnology, Worcester Polytechnic Institute, 100 Institute Road, Worcester, MA 01609, Fax: 508.

## Abstract

Bisphenol-A (BPA) is a ubiquitous precursor of polycarbonate plastics that is found in the blood and serum of >92% of Americans. While BPA has been well documented to act as a weak estrogen receptor (ER) agonist, its effects on cellular stress are unclear. Here, we demonstrate that high-dose BPA causes stress granules (SGs) in human cells. A common estrogen derivative, β-estradiol, does not trigger SGs, indicating the mechanism of SG induction is not via the ER pathway. We also tested other structurally related environmental contaminants including the common BPA substitutes BPS and BPF, the industrial chemical 4-nonylphenol (4-NP) and structurally related compounds 4-EP and 4-VP, and the pesticide 2,4-dichlorophenoxyacetic acid (2,4-D). The variable results from these related compounds suggest that structural homology is not a reliable predictor of the capacity of a compound to cause SGs. Also, we demonstrate that BPA acts primarily through the PERK pathway to generate canonical SGs. Finally, we show that chronic exposure to a low physiologically relevant dose of BPA disrupts SG assembly by inhibiting SGs upon additional acute stress. Our work identifies additional effects of BPA beyond endocrine disruption that may have consequences for human health.

## Introduction

Bisphenol A (BPA) is one of the highest volume chemicals produced globally, with an estimated 7.7 million metric tons produced in 2015 at an estimated value of US$15.6 billion (Research and Markets, 2016). BPA is used in the production of polycarbonate plastics and epoxy resins that are found in a myriad of consumer products including food packaging, receipts, automobiles, electronics, and medical devices. BPA enters the human body through ingestion or inhalation of plastic dust and particulates. A National Health and Nutrition Examination Survey (NHANES) study conducted by the Centers for Disease Control and Prevention (CDC) in 2003-2004 found that 92.6% of people tested had detectable levels of BPA in their urine (Calafat et al., 2008).

Many lines of evidence demonstrate the activity of BPA as an endocrine disrupting compound (Rochester, 2013, Ben-Jonathan, 2019). While the physiological relevance of this endocrine disruption is debated, many studies have linked BPA to infertility, obesity, diabetes, and cancer (Seachrist et al., 2016, Provvisiero et al., 2016, Qiu et al., 2016, (FDA), June 27, 2018, Apau J., 2018). The U.S. Food and Drug Administration (FDA) currently maintains that BPA is safe at the levels detected in the general population ((FDA), June 27, 2018). Still, public concern has led to pressure on manufacturers to remove BPA from consumer products. Most often, BPA is replaced with highly similar chemical analogs, such as BPS, BPF, or BPAF. Therefore, human exposure to the broader class of bisphenols is on the rise, and even less is known about the health effects of these BPA alternatives (Rosenmai et al., 2014).

An important consequence of BPA exposure that has been demonstrated in multiple contexts is oxidative stress, and the generation of reactive oxygen species (ROS)(Gassman, 2017). It is hypothesized that the metabolic breakdown of BPA may result in a greater cellular load of ROS than available antioxidants can manage, leading to oxidative damage (Gassman, 2017), and that these affects may be independent of the endocrine disrupting properties of BPA. For example, increased proliferation, ROS, and DNA damage were observed in cells lacking ER-α (Pfeifer et al., 2015). The exact mechanisms of BPA-related oxidative stress remain to be elucidated.

Stress granules (SGs) are cytoplasmic aggregates of proteins and mRNAs that form in response to stress-induced inhibition of mRNA translation (Anderson and Kedersha, 2009), and are biomarkers of the integrated stress response (Buchan et al., 2008, Farny et al., 2009, Kedersha et al., 1999). SGs are conserved among eukaryotic organisms from yeast to humans and are triggered by a wide variety of extracellular stresses including energy starvation, ion imbalance, oxidative stress, heat shock, and UV irradiation. The composition and dynamics of SGs vary based on the stressor (Aulas et al., 2017). SGs are generally thought to be protective, and conserve cellular resources by limiting mRNA translation during stressful circumstances. In some instances, SGs delay the onset of apoptosis in response to acute cellular stress, and the disruption of SGs can make cells more vulnerable to death (Arimoto et al., 2008). Indeed, the cellular consequences of endocrine disruption, such as oxidative stress, are also triggers for SG assembly, though direct links between endocrine disrupting compounds and SGs have yet to be examined.

SGs are a downstream consequence of a broader cellular stress response known as the integrated stress response. The integrated stress response is a mechanism by which a variety of environmental stresses (e.g., oxidative stress, UV irradiation, toxins) or biotic stresses (e.g., fever, viral infection) can signal the translation machinery to decrease protein synthesis, thereby conserving energy and permitting the cell to redirect resources to survival and the activation of various downstream stress response pathways (Pakos-Zebrucka et al., 2016). In mammalian cells, there are four kinases that can be activated during the integrated stress response: Heme-regulated eIF2α kinase (HRI), double-stranded RNA-dependent protein kinase (PKR), general control non-depressible kinase 2 (GCN2), and PKR-like endoplasmic reticulum kinase (PERK). Some stresses specifically activate one of the four kinases. For example, sodium arsenite specifically activates HRI (McEwen et al., 2005), and viral infection activates PKR (Garcia et al., 2006). These kinases once activated all target the same mechanism, the phosphorylation of serine 51 of the alpha-subunit of eIF2, which blocks the formation of initiation ternary complex (eIF2-GTP-tRNA^MET^) and thereby results in translational arrest. Often, the phosphorylation of eIF2α-Ser51 triggers the assembly of SGs (Kedersha et al., 2002), though SGs can also form through eIF2-independent mechanisms (Aulas et al., 2017, Farny et al., 2009). Whether BPA activates any of these kinase pathways was previously unknown.

Here, we investigated the link between the endocrine disrupting compound BPA and SGs. We determined that high doses of BPA cause canonical, phospho-eIF2α-dependent SG assembly via activation of PERK, one of four eIF2α kinases that mediate the integrated stress response. The common estrogen supplement β-estradiol does not trigger SG assembly, indicating that it is unlikely that BPA induced SG assembly in our cells is related to its reported endocrine disrupting activity. While BPA induces robust SG assembly, closely related BPA substitutes BPF and BPS cause lower or no SG response, respectively. Similarly, we find that the industrial chemical 4-nonylphenol (4-NP), but not highly related structural compounds 4-EP and 4-VP, trigger SG assembly. These results indicate that structural homology may not be a good predictor for environmental contaminant to promote SG assembly. Finally, we show that long term, low-dose BPA exposure within the physiologically relevant range can alter SG dynamics by reducing SG assembly upon later acute stress exposures, suggesting that real world chronic BPA exposures may compromise the ability of human cells to cope with environmental stress.

## Results

### High levels of BPA induce SGs but not processing bodies (P-bodies)

BPA has been associated with several potential health risks (Apau J., 2018, Seachrist et al., 2016, Qiu et al., 2016, Provvisiero et al., 2016). We wanted to test whether BPA triggers the integrated stress response using SG assembly as a biomarker. We treated cells with a range of BPA concentrations and exposure times, and used fluorescence microscopy to detect intracellular localization of the canonical SG component G3BP1. We determined that G3BP1-positive SGs form in ∼15% of U2OS cells after treatment with 300 µM BPA for 60 minutes (Fig. 1A-B). Treatment with 400 µM BPA for 60 minutes consistently induced granules in ∼100% of cells (Fig. 1A-B), therefore this concentration was used throughout our analysis.

**Figure 1:**
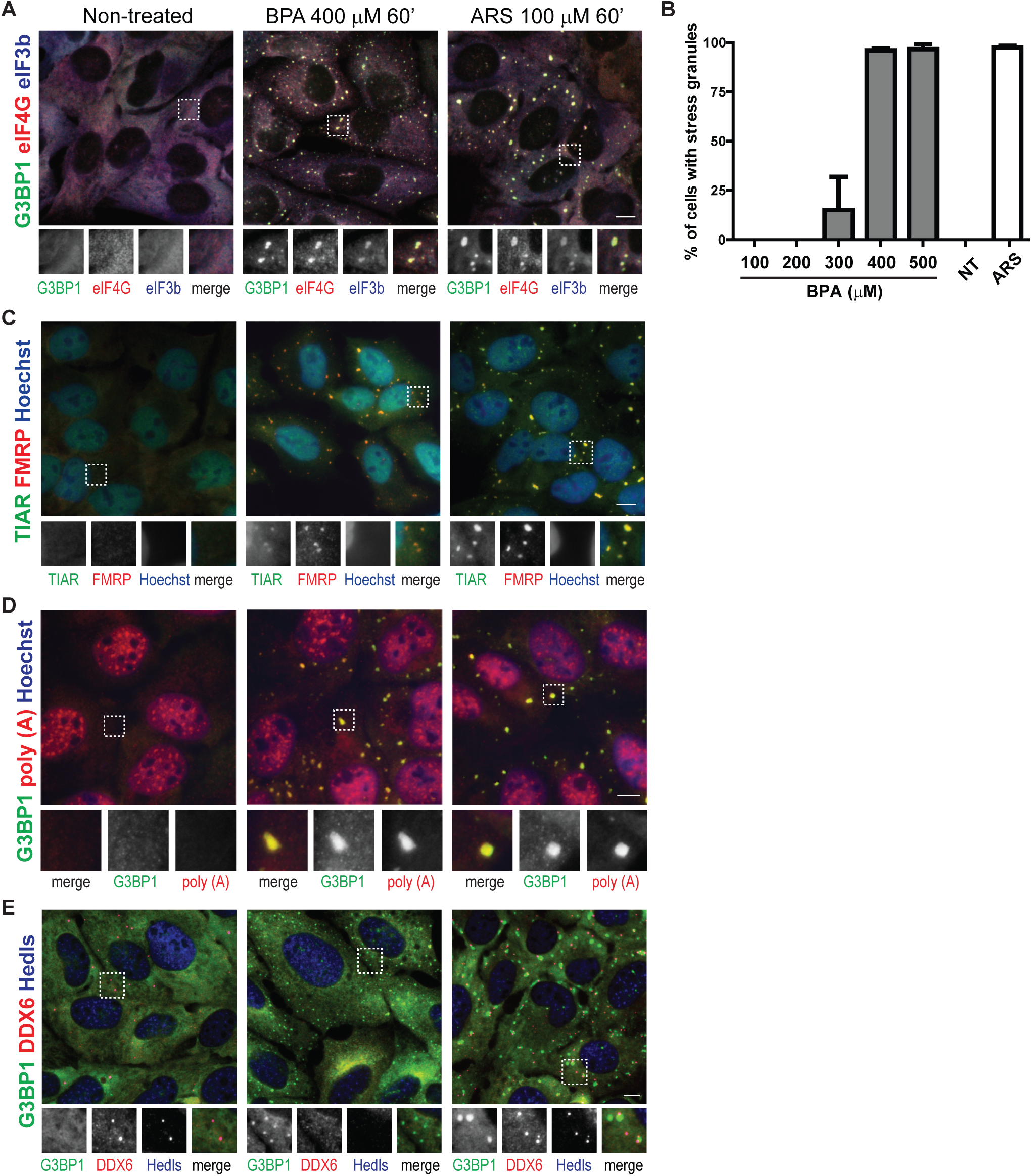
BPA promotes SG formation. A: Immunofluorescence of U2OS cells untreated (left) or treated with BPA (400 µM) (center) or arsenite (ARS, 100 µM) for 1 hour (right) detecting G3BP1 (green), eIF4G (red), eIF3b (blue). Boxed area is shown zoomed at 1.8X of original and individual channels are shown as black and white in order of green, red, blue then merged as a RGB image. B: U2OS cells were treated with the indicated range of BPA concentrations or 100 µM ARS for 1 hour, or left untreated (NT) then assessed for SG formation by immunofluorescence. Quantification of SGs was determined by the total number of cells containing two or more G3BP1 positive foci over the total number of cells. C: Same as A but detecting TIAR (green), FMRP (red), and Hoechst (blue). D: U2OS cells treated as indicated in B then fluorescence *in situ* hybridization to detect poly (A) RNAs (red) followed by immunofluorescence to detect G3BP1 (green) and counterstained with Hoechst (blue). E: Same as B but detecting G3BP1 (green), DDX6 (red), and Hedls (blue). Scale bar indicates 10 microns.

While G3BP is a widely-accepted marker for SGs, we employed other criteria to confirm that these BPA-induced G3BP foci were canonical SGs. To determine whether their composition was consistent with canonical SGs, cells were treated with BPA or sodium arsenite (ARS) as a positive control, then immunofluorescence was used to assess for colocalization of known SG markers with G3BP1. The translation initiation factors eIF4G and eIF3b colocalize with G3BP1 at cytoplasmic BPA-induced granules (Fig. 1A). The RNA binding proteins and classical SG markers fragile X mental retardation protein (FMRP) and T-cell intracellular antigen related protein (TIAR) colocalize in BPA-induced granules (Fig. 1C).

SGs contain polyadenylated (poly(A)) mRNA (Kedersha et al., 1999). We utilized fluorescence *in situ* hybridization (FISH) with an oligo dT probe to assess poly(A) RNA recruitment to BPA-induced granules. Poly(A) RNA consistently colocalizes with BPA-induced G3BP1-positive foci in a similar manner to ARS-induced SGs (Fig. 1D), indicating poly(A) mRNAs are localized to BPA-induced granules. Together these data indicate that high doses of BPA induce cytoplasmic foci that contain poly(A) RNA, G3BP1, FMRP, FXR1, eIF3b, eIF4G, and TIAR, which is consistent with BPA-inducing canonical SGs.

P-bodies are another cytoplasmic RNA granule assembled under some stress conditions (Kedersha et al., 2005). We assessed whether BPA also promotes P-body formation via immunofluorescence for the P-body markers Hedls and DDX6. We observed little to no focus formation of Hedls (also known as EDC4) or DDX6 in BPA treated cells while we did observe foci formation in ARS treated cells (Fig. 1E), indicating that unlike ARS, BPA does not promote P-body formation.

### High doses of BPA disrupt translation and promote SG condensation

Canonical SGs form upon translation initiation arrest and require G3BP1 and/or G3BP2 for their formation (Kedersha et al., 2016). To assess the effects of BPA on the translational status of cells, we utilized polysome profiling, a method of monitoring the relative translational state by density centrifugation, followed by measuring optical density at 254 nm across the eluted gradient. Treatment with SG-inducing levels of BPA caused collapse of polysome peaks, indicative of highly translated transcripts binding multiple ribosomes, and a corresponding increase in monosomes, or mRNAs that are initiated or being translated by one ribosome (Fig. 2A). This is consistent with the effect observed with SG-inducing levels of ARS (Fig. 2A).

**Figure 2:**
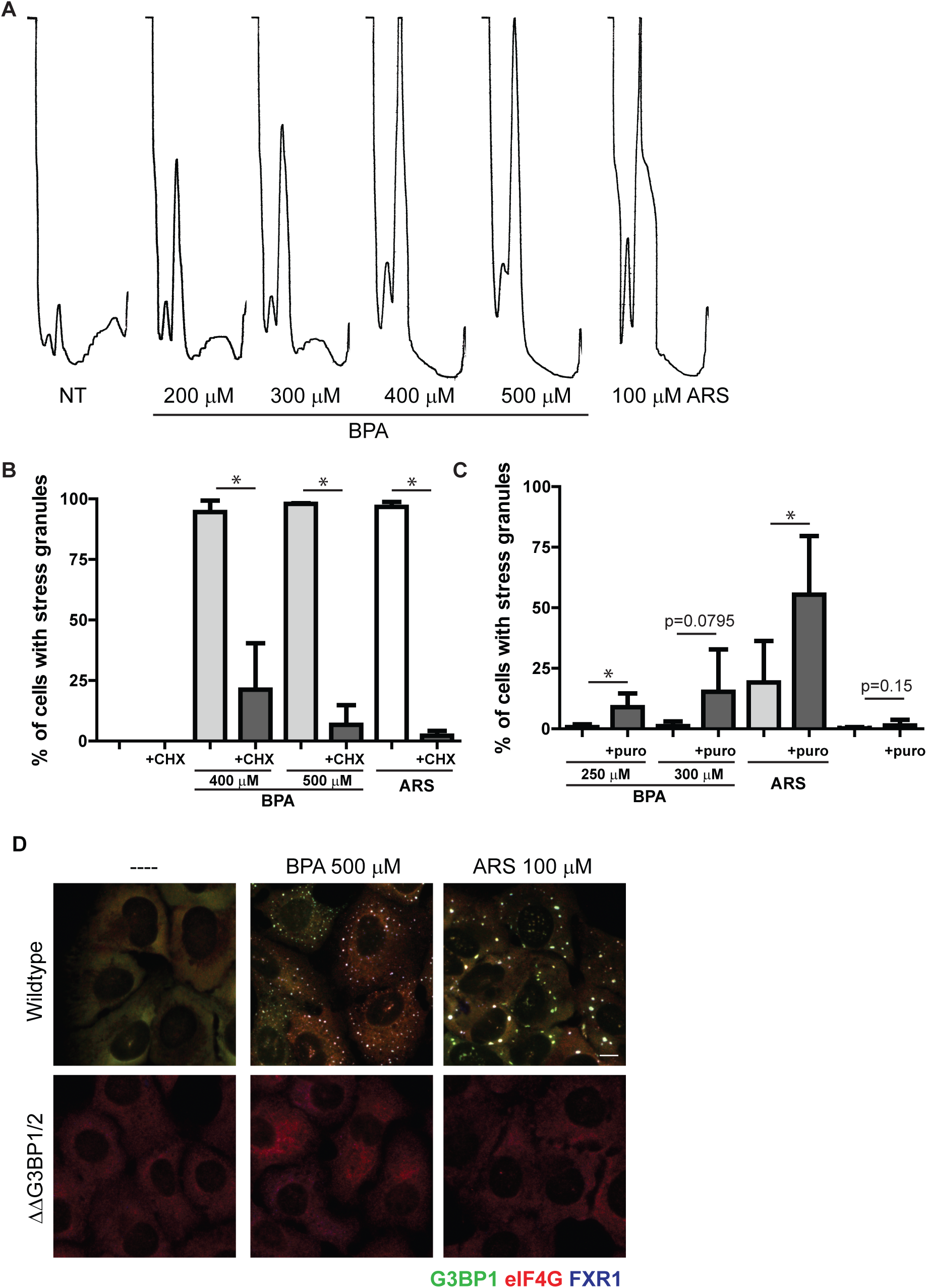
BPA induced SGs requires polysome disassembly and G3BP1/2 mediated condensation. A: Polysome profiles of U2OS cells treated with indicated concentrations of BPA or ARS for 60 minutes. Profiles were generated using OD 254 nm. B: U2OS cells were treated with BPA or ARS (100 µM) for 60 minutes. Where indicated, 50 µg/mL of cycloheximide was added for the final 30 minutes. Immunofluorescence was preformed and used to quantify the percentage of cells with SGs. Scale bar indicates 10 microns. C: U2OS cells were treated with indicate concentrations of BPA or ARS (50 µM) for 60 minutes, and cotreated with 20 µg/mL puromycin where indicated. Immunofluorescence analysis was used to quantify the percentage of cells with SGs. D: G3BP1 (green), eIF4G (red), and FXR1 (blue) in wildtype U2OS cells (top panel), or U2OS cells lacking G3BP1/2 (ΔΔG3BP1/2; bottom panel) treated with indicated concentration of BPA or ARS for 60 minutes or left untreated. B,C: Graphs represent mean with standard deviation, n≥3, * indicates 0.05≥p.

Canonical SGs are in dynamic equilibrium with polysomes. Experimentally, this can be assessed by blocking translocation of 40S subunits using the translation elongation inhibitor cycloheximide - which stabilizes polysomes and decreases SG formation, or and puromycin - which promotes premature ribosome termination and polysome disassembly, increasing dissociated into 40S subunits that are components of SGs and which promote SG formation (Kedersha et al., 2000). Cycloheximide treatment blocked BPA-induced SG assembly (Fig. 2B), and conversely, puromycin treatment promoted BPA-induced SG assembly at lower concentrations of BPA (Fig. 2C). Similar results were observed with the canonical SG inducer ARS (Fig. 2B, C).

The second step in SG assembly is condensation of SG components by the proteins G3BP1/2. G3BP1/2 are required for canonical SG formation downstream of polysome disassembly, and loss of both proteins prevents SG formation (Kedersha et al., 2016). To further characterize BPA-induced granules as canonical SGs, we treated SG-incompetent G3BP1/2 knockout U2OS cells with SG promoting levels of BPA, and confirmed that these cells lack the ability to form granules while wildtype U2OS cells form SGs (Fig. 2D, center panels). Similar results were observed with ARS (Fig. 2D, right panels). Together these data indicate that BPA induces translation initiation arrest and requires G3BP1/2 for the condensation of canonical SGs.

### PERK phosphorylation of eIF2α is required for BPA-induced SG assembly

Blocks in translation initiation can occur via several pathways, one of which includes phosphorylation of eIF2α by stress-activated kinases. In translation, eIF2α brings the initiator tRNA^Met^ to the AUG start codon, and phosphorylation of eIF2α halts global translation by preventing GTP/GDP exchange and activation of the eIF2/GTP/ tRNA^Met^ complex that recruits initiator tRNA^Met^ (Sonenberg and Hinnebusch, 2009). U2OS cells treated with a range of BPA concentrations displayed low eIF2α phosphorylation at 300 µM, increasing to robust levels at 400 µM, comparable to those induced by 100 µM ARS (Fig. 3A). This data is consistent with the concentrations required for SG assembly (Fig. 1A).

**Figure 3:**
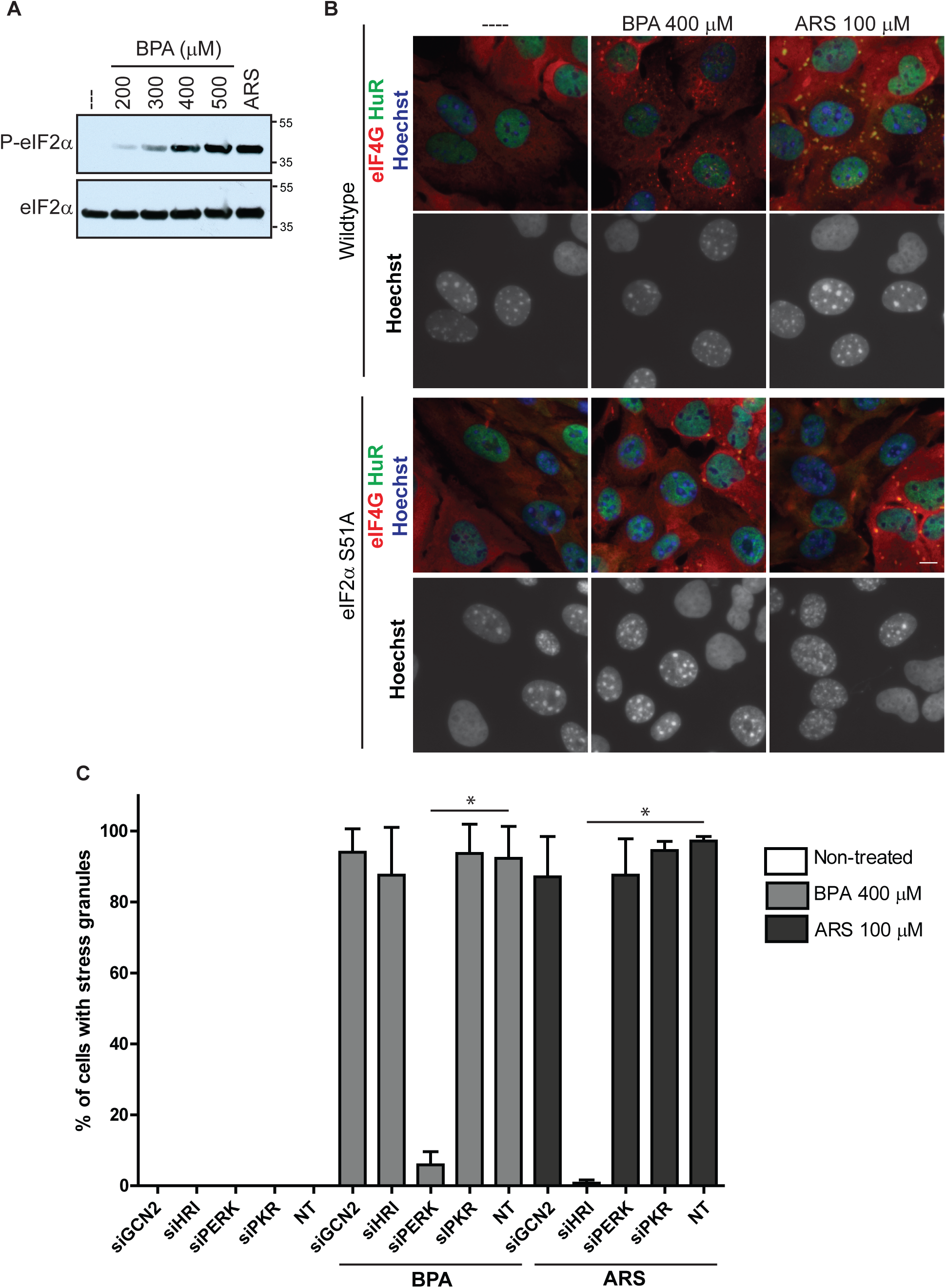
BPA-induced SG assembly requires PERK mediated phosphorylation of eIF2α. A: Immunoblot detecting levels of phosphorylated eIF2α (top blot, labeled P-eIF2α) and total eIF2α (bottom blot) in U2OS cells treated as indicated with BPA, ARS (100 µM) for 1 hour, or left untreated (indicated as ---). B: Immunofluorescence detecting HuR (green), eIF4G (red), and Hoechst/DNA (blue) in wildtype MEFs cocultured with U2OS cells (top panels), or MEFs with eIF2α-S51A point mutation cocultured with U2OS cells (bottom panels), treated as indicated with BPA or ARS for 1 hour or left untreated. MEFs exhibit punctate nuclear DNA. Scale bar indicates 10 microns. C: Quantification of SGs in U2OS cells following siRNA-mediated knockdown of GCN2, HRI, PERK, and PKR, prior to 1 hour BPA or ARS treatment, or no treatment. Graph represents mean with standard deviation, n≥3, * indicates 0.05≥p.

To assess whether eIF2α phosphorylation is required for BPA-induced SG assembly, we utilized mouse embryonic fibroblasts (MEFs) harboring a mutation that prevents eIF2α phosphorylation (eIF2α-S51A). Wildtype or eIF2α-S51A MEFs were co-plated with U2OS cells, which serve as an internal control, and then treated with SG-inducing levels of BPA or ARS. SGs were observed in wildtype MEFs and U2OS cells but not in eIF2α-S51A MEFs treated with BPA or ARS (Fig. 3B), indicating that eIF2α phosphorylation is required for BPA-induced SG formation.

To assess which eIF2α kinase is activated by SG-inducing levels of BPA, eIF2α kinases GCN2, HRI, PERK, or PKR were knocked down in U2OS cells with siRNA, then treated with SG-inducing levels of BPA or ARS as a control. Knockdown of GCN2, HRI, and PKR did not inhibit SG formation, but it was greatly decreased in cells treated with PERK siRNAs (Fig. 3C). Similarly, using a panel of human haploid (Hap1) cell lines with CRISPR/Cas9-mediated deletions of GCN2, HRI, PERK, or PKR (Aulas et al., 2017), we consistently found that cells lacking PERK failed to form SGs when treated with BPA (Fig. S1). Together these data indicate that BPA activates PERK to phosphorylate eIF2α and subsequently promote SGs assembly.

### BPF, 4-NP, and 2,4-D but not BPS, β-estradiol, 4-EP or 4-VP promote SGs

It is well-documented that BPA acts as an estrogen mimic and this contributes to its adverse effects (Rochester, 2013, Ben-Jonathan, 2019). β-estradiol (β-E) is an estrogen derivative that is generated from soy and often used in as a treatment for menopause. To assess whether β-E mimics BPA in promoting SGs, U2OS cells were treated with a range of β-E concentrations and assessed for SGs by immunofluorescence. Even at high concentrations β-E does not promote SG assembly (Fig. 4B). U2OS cells are not estrogen-responsive, and do not express ER-α or ER-β receptors (Tee et al., 2004), indicating that BPA likely does not work through an estrogen pathway to promote SG assembly in U2OS cells.

**Figure 4:**
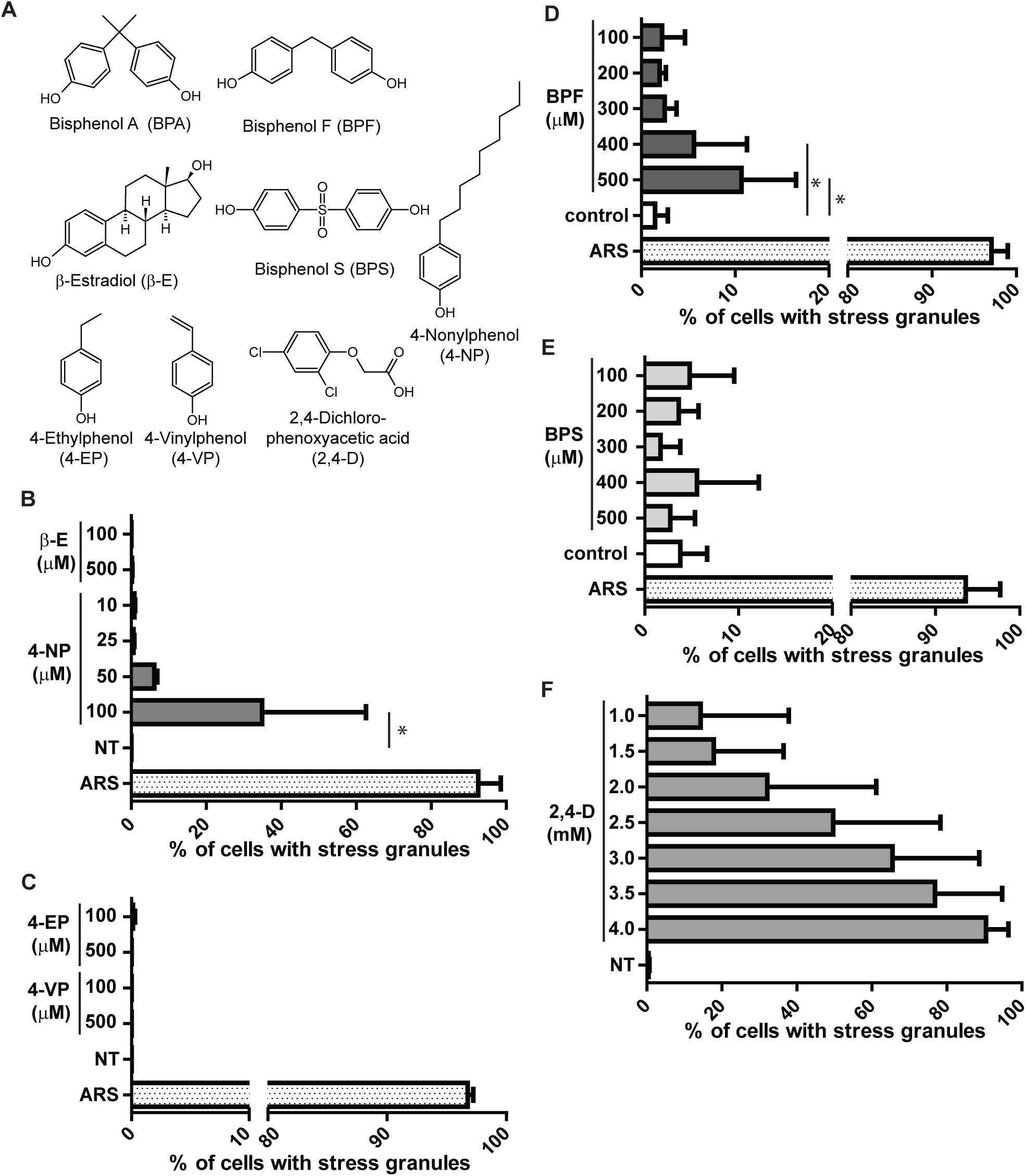
The BPA replacement BPF, the industrial chemical 4-NP, and the pesticide 2,4-D cause SG assembly. A: Molecular structures of chemicals used. Structures generated using ChemSketch. B: Quantification of SGs assessed by immunofluorescence in U2OS cells treated with indicated concentrations of β-estradiol (β-E), 4-nonylphenol (4-NP), or ARS for 1 hour or left untreated (NT). C: Same as B, but treated with 4-ethylphenol (4-EP), or 4-vinylphenol (4-VP). D-F : Quantification of SGs assessed by direct fluorescence GFP-G3BP1 in stably expressing U2OS cells treated with indicated concentrations of BPF, BPS, 2,4-dichlorophenoxyacetic acid (2,4-D), or ARS for 1 hour or mock treated (control). B-F: Graphs represent means with standard deviation, n≥3, * indicates 0.05≥p.

Like BPA, 4-nonylphenol (4-NP) is weakly estrogenic, used industrially in detergents and food packaging, and typically detected in human samples of urine and serum (Asimakopoulos and Thomaidis, 2015, Li et al., 2013, Calafat et al., 2005). We observed G3BP1, eIF4G, eIF3b positive granule form at 50-100 µM concentrations of 4-NP (Fig. 4B). Treating with higher doses of 4-NP promoted SG formation but also caused detachment and presumably death of U2OS cells (data not shown).

Structurally, 4-NP is a phenol ring with a nine carbon chain, and similar phenol derivatives are found naturally including 4-ethylphenol (4-EP) and 4-vinylphenol (4-VP). 4-EP and 4-VP differ from 4-NP in the length of their carbon tails (Fig. 4A). 4-EP and 4-VP are found as byproducts of bacterial metabolism and can be found at relatively high concentrations in beer and red wine (Langos and Granvogl, 2016, Teixeira et al., 2015). To test whether 4-EP or 4-VP promote SG assembly, U2OS cells were treated with a range of 4-EP and 4-VP concentrations and assessed for SG formation by immunofluorescence. Neither 4-EP nor 4-VP promoted SG formation at high concentrations (Fig. 4C), suggesting that longer carbon tails are required for SG formation.

Much public attention has been drawn to the potential adverse health effects associated with BPA, which has led to changes in BPA usage. Yet BPA is often replaced with bisphenol derivatives that remain largely untested (Rosenmai et al., 2014). We tested two commonly used BPA substitutes are bisphenol S (BPS) and bisphenol F (BPF). U2OS cells stably expressing GFP-G3BP1 were treated with a range of BPS or BPF then assessed by direct fluorescence for GFP-G3BP1 foci. Cells treated with BPF induced GFP-G3BP1 foci and quantification indicated considerably fewer cells assembled SGs compared to BPA (Fig. 4D) while BPS did not induce GFP-G3BP1 positive foci at the concentrations tested (Fig. 4E).

Lastly, we tested whether a commonly used pesticide, 2,4-dichlorophenoxyacetic acid (2,4-D) causes SG assembly. U2OS cells that stably express GFP-G3BP1 were treated with a range of 2,4-D concentrations and found to promote SGs between 1-4 mM concentration (Fig. 4F). Together these data indicate that like BPA, other environmentally relevant chemicals including 4-NP, BPF, and 2,4-D cause SG formation.

### Chronic low levels of BPA blunt the SG response

We have shown that high levels of BPA promote robust assembly of canonical SGs (Figs. 1-3), yet these high levels of BPA are rarely observed outside of industrial settings. Low levels of BPA are observed in most of the population (90-99% of individuals tested), ranging from 1-240 nM, with an average from 5-10 nM (Calafat et al., 2005). To assess how a chronic, low dose (referred to as chronic treatment) of BPA affects the cellular stress response, U2OS cells stably expressing GFP-G3BP1 were cultured in the presence of low BPA (5 nM) or left untreated for 24 hours then assessed for SG formation with SG inducing levels of BPA. When challenged with a SG-inducing level of BPA, cells that were treated chronically with BPA show a decrease in the ability to form SGs as compared to cells that were not treated with chronic BPA (Fig. 5B-C). It should be noted that chronic BPA treatment does not induce the formation of G3BP1 positive foci (Fig. 5B).

**Figure 5:**
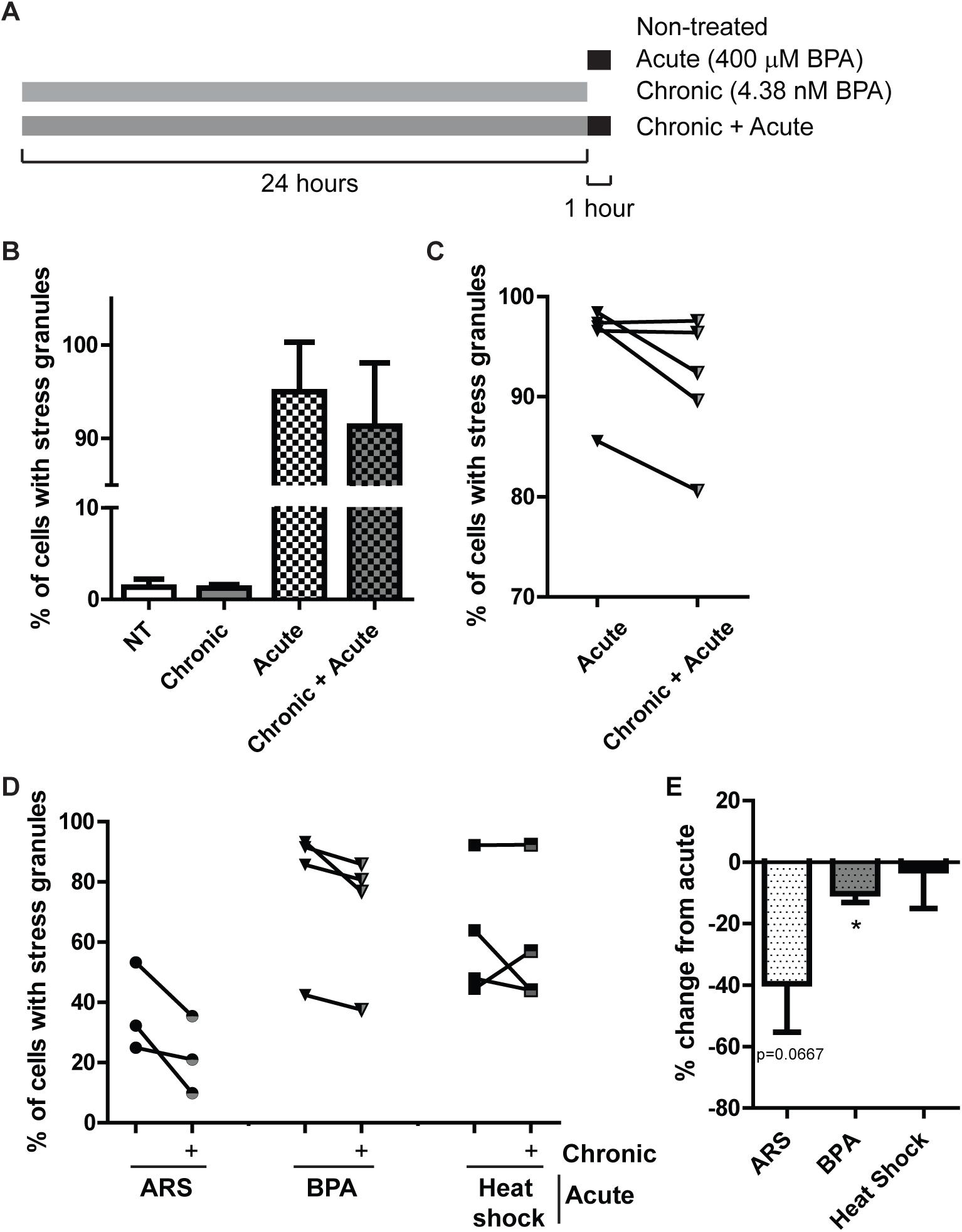
Chronic BPA exposure using physiologically relevant doses inhibits SGs assembly. A: Schematic indicating experiment performed to generate data in B-D. B: Quantification of SGs assess by direct fluorescence of GFP-G3BP1 in stably expressing U2OS cells treated with BPA by acute (400 µM, 60 minutes), chronic (4.38 nM, 24 hours) or both chronic and acute treatment. C: Same experiment (and data) as B showing how experimental samples pair as indicated by the line. Error bars in B and C represent +/- one standard deviation. D: Quantification of SGs detected by immunofluorescence in U2OS long term, chronically BPA treated (5 nM, 1 month; half grey half black shapes) or left untreated (solid black shapes) then treated with 75 µM ARS or 350 µM BPA for 60 minutes or heat shocked at 42°C for 25 minutes. Lines between chronically BPA treated and untreated samples indicate experimentally paired samples. E: Same experiment (and data) as D, but expressed as the percent change in SG formation between the acute-only treated sample and the paired chronic+acute treatment. Error bars and +/- standard error of the mean. n=3-5 replicates per condition, as indicated by paired data in C, D.

We then extended the duration of low BPA treatment to one month to assess long term chronic exposure. As seen with the 24 hour treatment, long term chronic treatment did not promote SGs (data not shown), yet challenging chronically treated BPA cells with arsenite or high-dose BPA dramatically reduced SG assembly (Fig. 5D). Calculating the percent change in SG formation between the acute-only treatments and the chronic-plus-acute treatments demonstrates a decrease in the ability of long term chronic BPA treated cells to form SGs when acutely challenged with arsenite or high-dose BPA, but not with heat shock (Fig. 5E). Together these data indicate that chronic treatment with low physiologically relevant levels of BPA can affect the cellular stress response by preventing SG assembly.

## Discussion

We show that high doses of BPA trigger the integrated stress response via activation of PERK to induce canonical SGs (Fig. 1-3). Besides BPA, we show that high levels of the BPA replacement BPF, the industrial chemical 4-NP, and the pesticide 2,4-D all cause SG assembly (Fig. 4). Together these results indicate that the integrated stress response is triggered by BPA, BPF, 4-NP, and 2,4-D, and indicate a need for more experimentation to define their health effects. Our work adds to a growing body of literature that highlights the physiological effect of BPA, and further indicates a need for caution in its application on everyday products.

Recent public outcry over the adverse health effects of BPA have caused the plastic industry to change its practices, and generate BPA-free products((FDA), June 27, 2018). Yet “BPA-free” products are often made using bisphenol derivatives such as BPF and BPS instead of BPA, and there is much less data on their health effects (Eladak et al., 2015). We observed that BPF, but not BPS, causes SG formation (Fig. 4), suggesting that the latter at least for the integrated stress response might be a less problematical substitute for BPA. In any case, caution should be used in replacing BPA with chemicals where less is known regarding their biological effects. Our data should not be over interpreted, and needs to be assessed in the context of additional studies.

Much BPA research has been centered on its potential endocrine disrupting properties. For example, in the recent CLARITY-BPA study, many of the study endpoints are endocrine-related properties such as ovarian and testicular development, animal growth, and fertility ((NTP), 2018). Our results show that while BPA induces SG assembly at high doses, while the estrogen mimic β-E does not induce SGs in U2OS cells (Fig. 4). Also tested here was the industrial compound and endocrine disrupter 4-NP, which we identified as promoting SG assembly (Fig. 4). U2OS cells do not express estrogen receptors ER-α or ER-β (Tee et al., 2004). While we cannot be certain that ER activation does not trigger SGs in other estrogen-responsive cell types, our data suggests that the mechanisms of BPA and 4-NP SG assembly are independent of their roles in endocrine disruption. Our findings indicate BPA triggers the integrated stress response via PERK activation, which aligns with other studies that show an oxidative stress effect of BPA (Gassman, 2017). Oxidative stress and altered SG dynamics have both been implicated in a range of disease states including neurodegenerative diseases (Wolozin, 2012), viral infections(Malinowska et al., 2016), and cancer (Anderson et al., 2015). Whether and how BPA contributes to oxidative stress and cellular damage in these diseases remains to be investigated.

BPA is thought to be quickly metabolized with a half-life of about five hours (Volkel et al., 2008, Volkel et al., 2002), and this short half-life could explain why BPA is observed in only 90% of patient samples. Interestingly, recent work has found BPA in sweat (Genuis et al., 2012), suggesting that BPA may accumulate in adipose tissue. In relation to our study, this could indicate that higher chronic levels could be observed, and that while BPA is considered to be quickly metabolized, it could also be stored and have high levels in particular tissues. We assessed the effects of chronic physiologically relevant, low nanomolar doses of BPA on cells (Fig. 5). We observed fewer SGs assembled when long term, low dose BPA treated cells were treated acutely with SG-inducing levels of BPA and ARS (Fig. 5). This data suggests that the preconditioning of cells with low doses of stress actually primes cells to respond differently. Altering the ability of a cell to assemble SGs could indicate the cell is primed to adapt to the stress, which may be a benefit to cell survival. Alternatively, it could indicate the cell has adapted and no longer considers a stress harmful, which could result in additional cell damage. Dampening of the integrated stress response could have a role in neurodegenerative diseases, cancer and viral infection, and potentially an altering the ability to form SGs has a role here. Alternatively, cells could be more adapted to the higher constitutive level of stress and mark that as their new baseline.

Our results suggest that structure alone is not a good indicator of the ability of a compound to induce a stress response (Fig. 4). It is also important to consider that humans currently have contact with as many as 85,000 chemicals obtained from their environments (EPA) some of which may be having toxic effects at their typical exposure levels (Sturla et al., 2014), others of which may be inducing protective effects (Calabrese and Mattson, 2017). It will therefore be critical to continue to examine not only the range of effects of these individual substances, but also the compound effects of substances within the complex chemical context that human cells currently reside. For example, as BPA loses favor with the manufacturing industry due to social and political pressure, the use of BPS, BPF and other chemical analogues will increase. There is already preliminary evidence that bisphenols and other related compounds have synergistic effects that can affect the cellular consequences of exposure (Kortenkamp, 2007, Le Magueresse-Battistoni et al., 2018). The policy implications of this information are profound: it will not be sufficient to continue to make one-off bans on individual chemicals, only to have them replaced by others about which even less is known. The future of ensuring safer environments will depend upon a more holistic understanding of the effects of chemical classes on cellular physiology and human health.

## Material and Methods

### Cell culture and drug treatments

Cells (U2OS, MEF with or without eIF2α-S51A, Hap1 cells) were maintained at 37°C in a CO2 incubator in DMEM (Corning) supplemented with 10% FBS, 20 mM Hepes (Gibco), 1% penicillin/streptomycin. For SG induction, cells were treated with chemicals purchased from Sigma including Bisphenol A (BPA), Bisphenol S (BPS) Bisphenol F (BPF), 4-nonylphenol (4-NP), 4-ethylphenol (4-EP), 4-vinylphenol (4-VP), β-estradiol (β-E), sodium arsenite (ARS), and 2,4-dichlorophenoxyacetic acid (2,4-D). As indicated in figure legends, cycloheximide (CHX) and puromycin (puro) were added. For chronic treatments, BPA (5 nM final concentration from 5 µM stock) or mock treatment (methanol) was added when cells were split at the same time every 2-3 days at ∼90% confluency. For 1 month BPA or mock treatments, each n represents a separate flask of cells split.

### Polysome profiling

5×10^5^ cells were plated onto 6 well dishes and the following day treated as indicated. Then cells were treated with 100 µg/mL cycloheximide for 5 min, washed once with HBSS, and 500 µL of polysome buffer (20 mM Tris, pH 7.4, 150 mM NaCl, 5 mM MgCl2, 1% Tritonx100, RNaseIN, 1 mM DTT, 100 µg/mL cycloheximide) was added. Cells were scraped and collected into 1.7 mL epi tubes and rotated at 4°C for 10 min and centrifuged at 10,000 x g for 10 min at 4°C. The supernatant was loaded onto a preformed 10-50% sucrose gradient made in polysome buffer. Gradients were centrifuged in a Beckman Sw55Ti at 45,000 rpm for 100 min at 4°C. Then a Brandel bottom-piercing apparatus connected to an ISCO UV monitor and pump extracted the gradient while measuring the eluate at 254 nm.

### Immunofluorescence

1×10^5^ U2OS cells were plated into a 24 well dish seed with coverslips. The following day cells were treated as indicated in figure legends. Then the cells were fixed with 4% paraformaldehyde for 15 min, permeabilized with −20°C methanol for 5 min, and blocked for at least 20 min with 5% normal horse serum (NHS; ThermoFisher) diluted in PBS. Primary antibodies were diluted in blocking solution and incubated for 1 hour at room temperature or overnight at 4°C. Antibodies against G3BP1(sc-365338), eIF4G (sc-11373), eIF3b (sc-16377), FMRP (sc-101048) FXR1 (sc-10554), TIAR (sc-1749), Hedls (sc-8418) and HuR(sc-5261) were all purchased from Santa Cruz and used at a 1:250 dilution. The DDX6 antibody was purchased from Bethyl Laboratories (A300-461A) and used at a 1:500 dilution. Coverslips were washed 3 times for 5 min then incubated with secondary antibodies and Hoechst 33258 (Sigma) for 1 hour at room temperature and again washed. Coverslips were mounted on glass slides with Vinol.

### Direct fluorescence

7×10^4^ U2OS cells stably expressing GFP-G3BP1, RFP-DCP1A (REF) were plated in 12-well plates seeded with coverslips and two days later were treated as indicated. Cells were fixed with 4% paraformaldehyde for 10 minutes, permeabilized with ice cold methanol for 10 minutes, and rinsed with PBS. Hoechst 33258 (Sigma) was used to stain nuclei. Coverslips were mounted on glass slides with Vinol.

### Microscopy

Wide-field fluorescence microscopy was performed using an Eclipse E800 microscope (Nikon) equipped with epifluorescence optics and a digital camera (Spot Pursuit USB) or an AXIO observer A1 (Zeiss) equipped with epifluorescence optics and a digital camera (SPOT Idea 5.0mp). Image acquisition was done with a 40X air or 60X oil objective. Images were merged using Adobe Photoshop.

### siRNA treatment

1.8×10^5^ U2OS cells were seeded in the 6-well plates and grown for 2 hours, then, transfected using100 pmol siRNA (SmartPool-Dharmacon, Thermo Scientific), Lipofectamine 2000 (Invitrogen) in OPTI-MEM (Life Technologies). Cells were incubated with siRNA for 24 hours at which time a second transfection was done under the same conditions and 24 hours later the cells were collected, counted, and plated for experiments.

### Fluorescence *in situ* hybridization (FISH)

U2OS cells were treated as indicated in figure legends then fixed with 4% paraformaldehyde for 15 minutes, permeabilized with −20°C methanol for 10 minutes, then dehydrated in 70% ethanol at 4°C. Cells were stored at 4°C for at least overnight then wash several times with 2X saline sodium citrate (SSC; Ambion). Cell were equilibrated in hybridization buffer (Sigma) for 15 minutes at 65°C then incubated with oligo dT40X probe for 1 hour at 42°C. Cells were washed twice with 2X SSC for 10 minutes at 42°C then several times at room temperature. Coverslips were then processed for immunofluorescences as indicated above (from block step on).

### SG quantification

At least three independent 40X images from at least three independent experiments were counted as the total SG positive over the total number of cells. Cells were counted as SG positive if at least two G3BP1 foci were present in the cytoplasm. The total number of cells was quantified by counting nuclei using Hoechst. Graphs were generated with GraphPad Prism and error bars indicate standard deviation.

### Immunoblotting

5×10^5^ U2OS cells were plated in 6 well dishes and the following day drug treated as indicated. Cells were washed once with PBS then lysis buffer (50 mM Hepes pH 7.6, 5% glycerol, 150 mM NaCl, 0.5% NP40, 1X Halt protease inhibitors (ThermoScientific), 1X Halt phosphatase inhibitors (ThermoScientific)) and cells were scraped and collected into tubes. Cells were sonicated using a cup sonicator and the protein concentration was determined using BioRad Protein Assay (BioRad). SDS dye was added to a 1X concentration and lysates (equal µg of protein) were loaded onto 4-20% Tris-glycine gels (BioRad). Gels were transferred to nitrocellulose, then blocked with 5% milk in TBST. Primary antibodies were diluted in 5% NHS and incubated overnight at 4°C. p-eIF2α (Abcam, ab32157), eIF2α (Cell Signaling, 2103). Secondary antibodies were diluted 1:10,000 in 5% NHS and incubated for 1 hour at room temperature. Blots were washed at least three times for 10 min after antibody incubation. HRP signal detected using SuperSignal West Pico Chemiluminescent Substrate (Thermo Scientific),

### Molecular structures

Molecular structures were generated with ChemSketch by ACD Labs Freeware 2016.

### Statistical Analysis

Statistical analysis of an unpaired t-test was preformed where indicated and statistical significance was considered p<0.05. n indicates the number of replicate experiments and within each experiment and at least 100 cells were counted.

---

**Figure S1:**
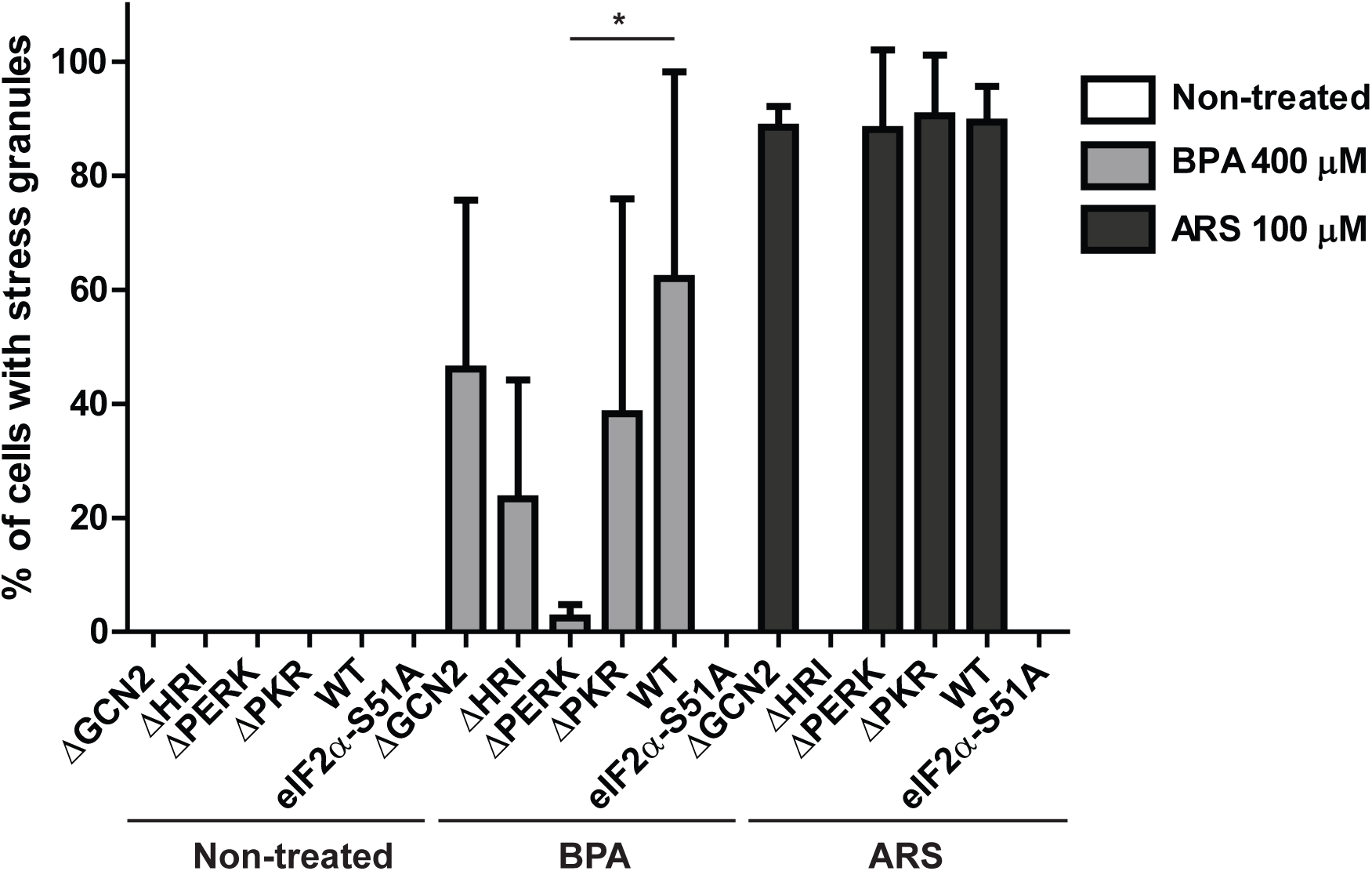
PERK promotes eIF2α-S51A phosphorylation in response to BPA. Quantification of SGs detected by immunofluorescence in Hap1 cells that were genetically modified with CRISPR to knockout GCN2, HRI, PERK, PKR, or to contain the point mutation eIF2α-S51A. Cells were treated with BPA or ARS for 1 hour, or left untreated. Graph represent means with standard deviation, n=3, * indicates p<0.05.

